# mRNA 5′ terminal sequences drive 200-fold differences in expression through effects on synthesis, translation and decay

**DOI:** 10.1101/2022.02.17.480814

**Authors:** Antonia M.G. van den Elzen, Maegan J. Watson, Carson C. Thoreen

## Abstract

mRNA regulatory sequences control gene expression at multiple levels including translation initiation and mRNA decay. The 5′ terminal sequences of mRNAs have unique regulatory potential because of their proximity to key post-transcriptional regulators. Here we have systematically probed the function of 5′ terminal sequences in gene expression in human cells. Using a library of reporter mRNAs initiating with all possible 7-mer sequences at their 5′ ends, we find an unexpected impact on transcription that underlies 200-fold differences in mRNA expression. Library sequences that promote high levels of transcription mirrored those found in native mRNAs and define two basic classes with similarities to classic Initiator (Inr) and TCT core promoter motifs. By comparing transcription, translation and decay rates, we identify sequences that are optimized for both efficient transcription and growth-regulated translation and stability, including variants of terminal oligopyrimidine (TOP) motifs. We further show that 5′ sequences of endogenous mRNAs are enriched for multi-functional TCT/TOP hybrid sequences. Together, our results reveal how 5′ sequences define two general classes of mRNAs with distinct growth-responsive profiles of expression across synthesis, translation and decay.

## Introduction

Genes are expressed in complex patterns that are post-transcriptionally controlled through instructions encoded within mRNAs. These instructions can occur throughout mRNAs, but 5′ terminal sequences have unique regulatory potential because they are adjacent to the mRNA cap structure. This structure is appended to mRNAs co-transcriptionally and then rapidly bound by nuclear cap-binding proteins that impact both splicing and export from the nucleus [1]. Once in the cytoplasm, the cap is bound by a different complement of factors that facilitate translation [1]. Finally, the removal of the cap is the penultimate and generally irreversible step of mRNA decay.

There are several known examples of 5′ terminal sequences with post-transcriptional regulatory functions. The best-studied of these are terminal oligopyrimidine (TOP) motifs, which are defined as a +1 C followed by a series of pyrimidines [2]. These motifs are common on mRNAs encoding translation factors and render their translation and stability sensitive to growth signals through the mTOR Complex 1 pathway [3–6]. Regulation of these mRNAs is mediated through the translation repressor La-related protein 1 (Larp1), which binds TOP motifs and sequesters the mRNAs away from the cap-binding translation factor, eIF4F [7–9]. eIF4F also interacts directly with cap-adjacent nucleotides [10] and has intrinsically lower affinity for mRNAs initiating with a +1 C [11]. Finally, there is evidence that decapping efficiency is influenced by 5′ nucleotides, as in yeast the decapping protein Dcp2 preferentially cleaves mRNAs with A or G as the +1 nucleotide, potentially reducing the overall stability of these transcripts [12].

5′ terminal sequences are also constrained by requirements for efficient transcription start site selection and initiation. These sequence patterns are reflected in the “core promoter” sequence [13], a 100-150 nt region that encodes an array of elements that position the transcription machinery and ultimately trigger transcription initiation. There is no canonical sequence that defines the transcription start site (TSS), and transcription generally initiates at a spectrum of positions. Nonetheless, the most common TSS motif is the Initiator (Inr) sequence, which occurs in ∼50% of genes [13]. The human Inr consensus sequence is BBCA_+1_BW [14] (B=G,C,T; W=A,T; +1 indicates the first nucleotide), although a global analysis of TSS data reported a minimal Inr of YR_+1_ [15, 16]. A less common initiation motif is the TCT motif [17].

Similarly to TOP motifs, this element is most common in the promoters of translation-associated genes. The established motif, based primarily on analyses of endogenous RP mRNA TSSs, is defined as YYC_+1_TTTYY, yielding mRNAs that initiate with a +1 C rather than the +1 A that characterizes Inr TSSs [17].

Aside from these examples, the full spectrum of 5′ sequence functions has not been determined. A powerful approach for identifying similar regulatory elements has been to use large-scale reporter libraries that systematically query the function of thousands of mRNA sequences [18–22]. However, these systems are generally designed with promoters that produce mRNAs with fixed 5′ terminal sequences, and so have not assessed functions of different 5′ sequences. To survey functions of 5′ sequences directly, we constructed a reporter library of mRNAs that vary only in the first 7 5′ nucleotides. Library mRNAs were unexpectedly expressed at a wide range of levels that closely parallel observations for endogenous mRNAs and share features with Inr and TCT motifs. The translation and stability of library mRNAs is similar under basal conditions but diverge under conditions when cap-dependent translation is repressed. The 5′ sequences that define translationally-regulated mRNAs include TOP motifs and related sequences that partially overlap with TCT sequences that are optimal for transcription. We show that endogenous mRNAs are enriched for hybrid TOP/TCT sequences that combine the efficient transcription of TCT motifs and the translation regulation of TOP sequences, and that these patterns are evolutionarily conserved.

## Results

### Design of a 5′ reporter system

To systematically test the functions of 5′ terminal sequences, we constructed a plasmid encoding a CMV promoter, a short 5′ UTR, a coding sequence (Renilla luciferase), and a constant 3′ UTR (Figure 1A). The CMV promoter preferentially initiates transcription at a specific site [9], allowing us to specify the 5′ ends of expressed mRNAs. A cassette of 7 random nucleotides was then positioned directly after the CMV promoter region. The rationale for choosing a 7 nt sequence was that many RNA-binding proteins interact with similar size sequences and it is short enough that all 16,384 7 mers could be robustly quantified by deep sequencing. Deep-sequencing of the plasmid library identified 16359 of 16384 7-mer sequences with greater than 20 reads and no position-specific bias (Supplemental Figure 1A). There is a slight overrepresentation of T and G nucleotides, which are likely artifacts introduced during the initial synthesis of the randomized 7 nt sequence. This minor bias was corrected for in downstream analysis (Methods). This plasmid library, which we refer to as the 5pseq library, was then packaged into lentivirus and used to stably infect HeLa cells.

**Figure 1.**
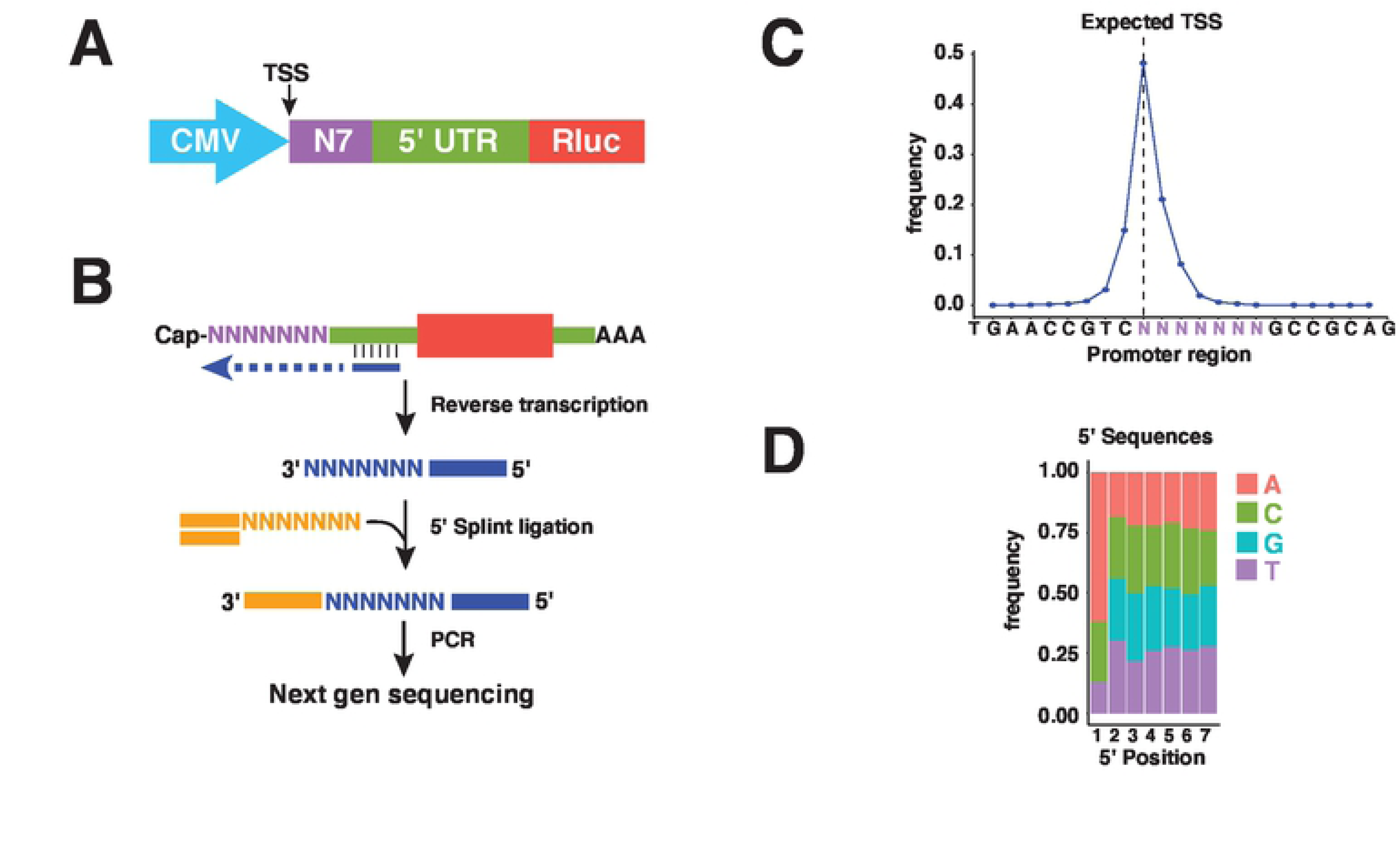
5pseq library design. (A) Organization of the 5pseq expression plasmid. (B) Strategy for library preparation. Extracted mRNA is reverse transcribed with a library-specific primer. A 5′ linker is then attached by splint-ligation, followed by PCR amplification with Illumina-compatible primers. (C) 5pseq mRNAs initiate at the expected TSS. Library mRNAs expressed in HeLa cells were processed as in (B) and analyzed to determine initiation sites within the 5pseq promoter region (excluding +1 G mRNAs). (D) Nucleotide frequencies at the 5′ ends of 5pseq mRNAs. Library mRNAs expressed in HeLa cells were processed as in (B) and used to determine the frequency of all 5′ 7-mer sequences initiating with A, C, or U. Frequencies of each 7-mer were then normalized to frequencies in the plasmid library and used to determine the frequency of each nucleotide at each position.

We next measured the expression levels of all library mRNAs. RNA was extracted from the infected HeLa cells, reverse transcribed with a library-specific primer, and then appended with a 5′ linker using splint-ligation (Figure 1B). Sequencing libraries were amplified and analyzed by deep sequencing, confirming preferential initiation at a single position (∼48%) (Figure 1C). A downside of this approach is that commonly available reverse transcriptases often append a non-templated cytidine to the 3′ end of the synthesized cDNA [23]. This appears as an additional 5′ G in sequencing libraries. This addition was corrected computationally for library mRNAs initiating with A, C or U. However, we could not distinguish between templated and non-templated +1 G nucleotides, and so excluded these sequences from further analysis, leaving 12,049 of 12,288 possible sequences with +1 A, C or U detected with greater than 50 reads in all replicates. Aside from the +1 position, nucleotides were observed at roughly the expected frequencies (Figure 1D).

### Library mRNAs are expressed across a 200-fold range

Unexpectedly, the expression levels of library mRNAs spanned more than a 200-fold range and followed a distinctly bimodal distribution (Figure 2A). Inspection of the underlying sequences revealed that high and low-expressed mRNAs showed significantly different preferences for the first (+1) nucleotide (Figure 2A). mRNAs initiating with an ‘A’ were uniformly well-expressed, while those initiating at a ‘U’ were poorly expressed. In contrast, mRNAs initiating with a +1 C nearly spanned the entire range (Figure 2A). Amongst the 10% most highly-expressed +1 A motifs, there was a notable depletion of A nucleotides throughout the remainder of the 7 nt sequence. The top 10% +1 C motifs also showed a decreased frequency of A nucleotides throughout the 7 nt sequence, but additionally required a +2 T/U for efficient transcription (Figure 2B). To test the importance of the +2 position for these mRNAs, we generated a series of reporters with different +2 nucleotides. Expression analysis of these reporters confirmed the importance of a T/U for maximal expression (Figure 2C). These results argue that TSS sequences have a major impact on mRNA expression level.

**Figure 2.**
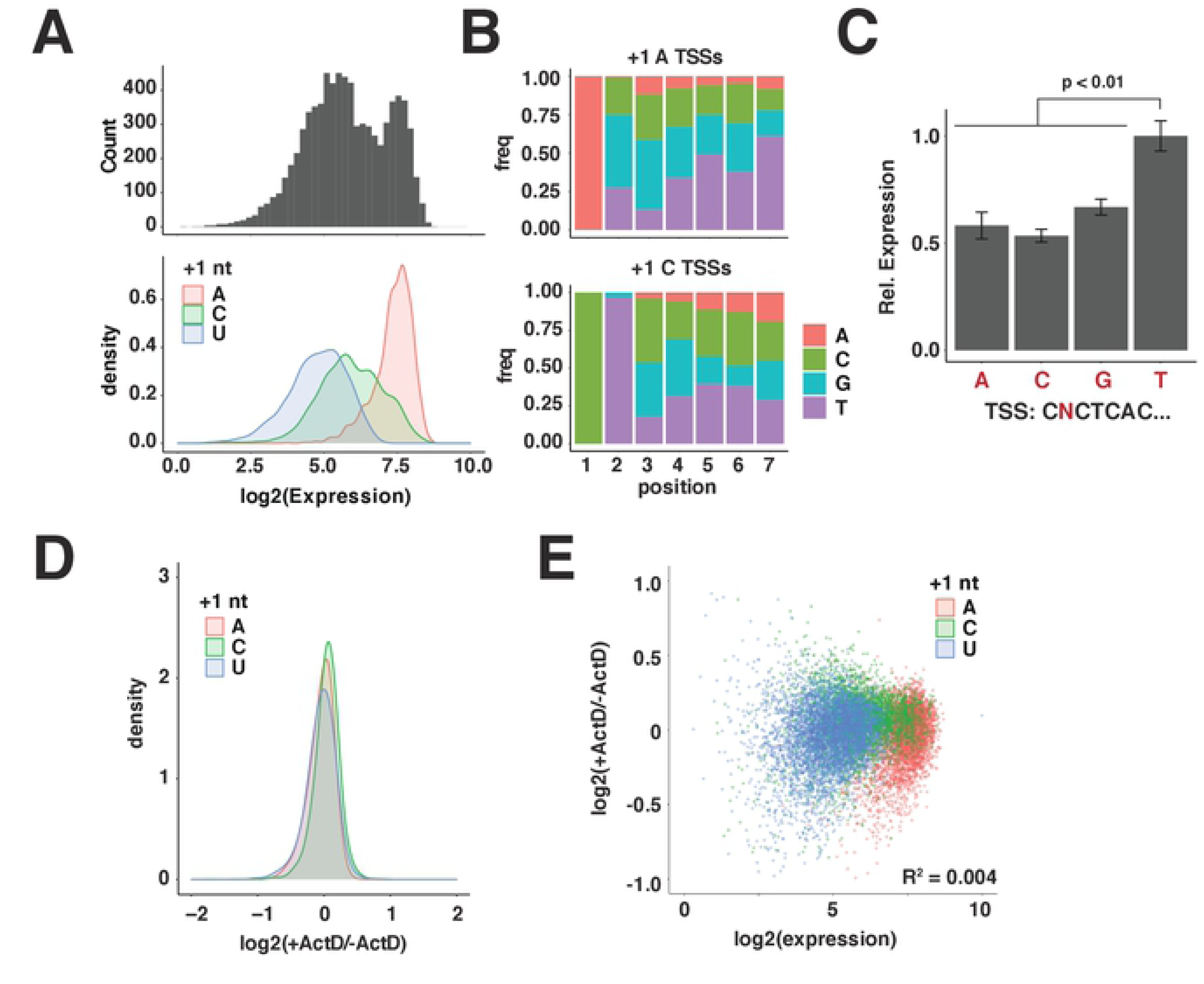
mRNA 5′ sequences strongly impact expression level. **(A)** Expression levels of 5pseq library mRNAs. Library mRNAs were expressed in HeLa cells, extracted, and used to prepare sequencing libraries. Frequencies of all 7 nt sequences initiating with A, C or U were determined and normalized to frequencies within the input plasmid library. Library mRNAs were then binned by expression level and plotted as a histogram (upper panel) or density plot separately depicting expression levels of mRNAs initiating with the indicated nucleotide (lower panel). **(B)** Motifs in 5′ sequences of well-expressed +1 A and +1 C mRNAs. Nucleotide frequencies within the first 7 nucleotides of the top 5% +1 A and +1 C library mRNAs from (A). **(C)** The +2 U is required for maximal expression of +1 C mRNAs. Plasmids expressing mRNAs encoding Renilla luciferase and initiating with the indicated 5′ sequences were transfected into HEK-293T cells. Expression levels of each mRNA were determined 24 h later by qPCR. Significance by t-test between +2 U and each other construct, n=3. **(D)** Decay rates of library mRNAs are similar under control conditions. HeLa cells expressing library mRNAs were treated with 2 µg/mL Actinomycin D for 0 or 2 h in 2 biological replicates. Libraries prepared from extracted mRNA were analyzed to determine relative changes in levels between ActD-treated (ActD+) and untreated (ActD-) conditions. Log2 changes in levels are plotted separately for mRNAs initiating with the indicated nucleotide. **(E)** Expression levels of library mRNAs are not correlated with decay rates. Decay rates (ActD+/ActD-) from (D) are compared with expression levels of library mRNAs from (A) expressed in HeLa cells.

Expression levels are determined by both synthesis and decay rates. To determine differences in stability, we measured changes in expression levels following transcription shutoff with the Pol II inhibitor Actinomycin D. Under these conditions, the relative decay rates of nearly all 5′ sequences fell within a 2-fold range (Figure 2D) and showed minimal correlation with expression level (Figure 2E). We therefore concluded that the large range in expression levels of mRNA primarily reflects differences in transcription rates.

### Library 5′ sequence expression predicts the expression of endogenous 5′ sequences

Next, we examined the potential significance of these findings for understanding differences in expression levels between endogenous mRNAs. To test this, we analyzed several previously reported TSS datasets prepared using cap-analysis of gene expression (CAGE), a strategy for determining mRNA 5′ terminal sequences by isolating and then sequencing RNA fragments with 5′ caps [24]. For each dataset, we selected reads that aligned to “promoter regions” of genes, which we defined as a 1 kb window surrounding the annotated transcription start site for each transcript. We then compared the frequencies of 5′ 3mer sequences, reasoning that these account for most of the variation in expression observed in the 5pseq library. Strikingly, the relative expression of 3mer TSS sequences in each of these CAGE datasets was significantly correlated with expression level in the 5pseq library (Figure 3A). In both CAGE and 5pseq datasets, +1 A mRNAs were collectively well-expressed, +1 U mRNAs were poorly expressed, and +1 C mRNAs spanned a broad range. The similarity between results from library and endogenous mRNAs argue that 5′ terminal sequences are important determinants of expression level.

**Figure 3.**
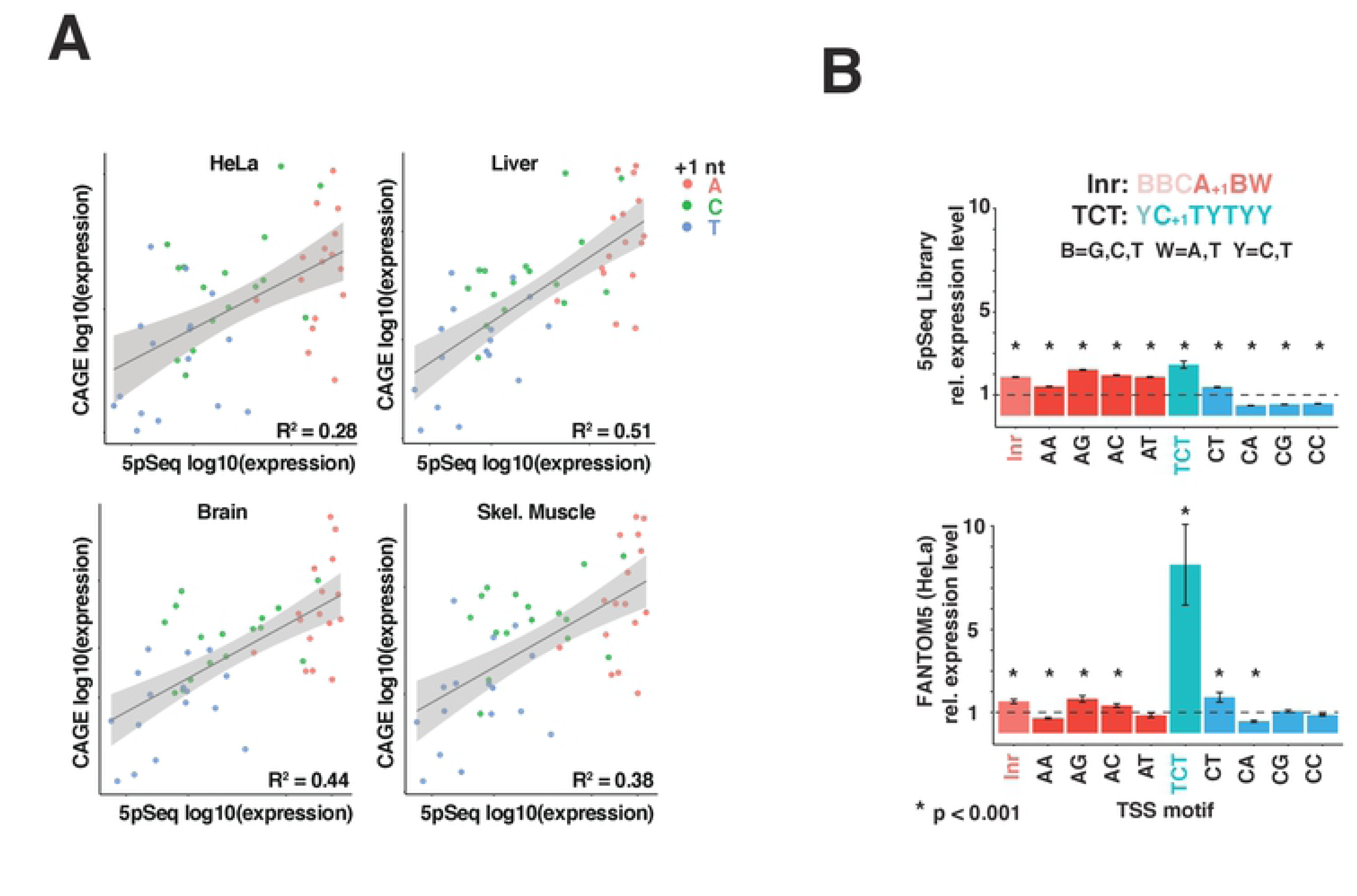
Expression levels of library TSSs correlate with endogenous TSSs. **(A)** Comparison of library and endogenous TSS expression. CAGE reads from HeLa cells and the indicated human tissues were extracted from the 1 kb region surrounding the 5′ ends of annotated transcripts and used to determine the frequencies of 5′ 3-mer sequences. CAGE 3mer frequencies were then compared to library 3mer frequencies from Figure 2A. Significance level of p<10^-5^ for all plots. **(B)** AG and CT are preferred TSS sequences in library and endogenous mRNAs. Expression levels of the indicated di-nucleotide motifs were compared to classical Inr and TCT sequences for library and endogenous TSSs. Significance determined by t-test, compared to mean expression of all TSS sequences (indicated as a dashed line).

Previous analyses of endogenous promoters have identified Inr and TCT motifs as important determinants of TSS selection [25]. These motifs include nucleotides upstream of the TSS but yield mRNAs with specific 5′ sequences. We therefore wondered whether these motifs are also linked to expression level in library mRNAs. Indeed, both Inr and TCT classes of 5′ sequences (ABW (B=C,G,T; W=A,T) for Inr and CTYTYY (Y=C,T) for TCT motifs) were expressed at significantly higher levels in both library and endogenous mRNAs (Figure 3B). Given the key role of the first two nucleotides in the expression of library mRNAs, we wondered whether these are the key predictive features of Inr and TCT motifs. Both AG and CT 5′ sequences were indeed expressed at significantly higher levels than average (Figure 3B). AG, in particular, was expressed at even higher levels than Inr sequences in both datasets, suggesting that this dinucleotide sequence is an optimal version of the Inr, consistent with previous analysis of CAGE data [15, 16]. In contrast, the classical TCT 5′ sequence was a much stronger predictor of expression in CAGE data than in 5pseq library mRNAs (Figure 3B). This suggests that endogenous TCT sequences might be also optimized for functions beyond efficient transcription initiation, such as post-transcriptional control.

### 5′ sequences define distinct patterns of translation

To determine post-transcriptional functions of 5′ terminal sequences, we first examined translation. Towards this end, extracts from HeLa cells stably expressing the 5pseq library were separated into sub-polysome and polysome-associated fractions by centrifugation through sucrose gradients (Figure 4A). Translation levels were estimated by calculating polysome/sub-polysome (P/SP) ratios for each library mRNA. Under basal conditions, library mRNAs showed small differences in translation rates, which varied across a 4-fold range (Figure 4B). mRNAs were similarly well translated regardless of the +1 nucleotide, although +1 A mRNAs were translated slightly better than +1 C or U mRNAs.

**Figure 4.**
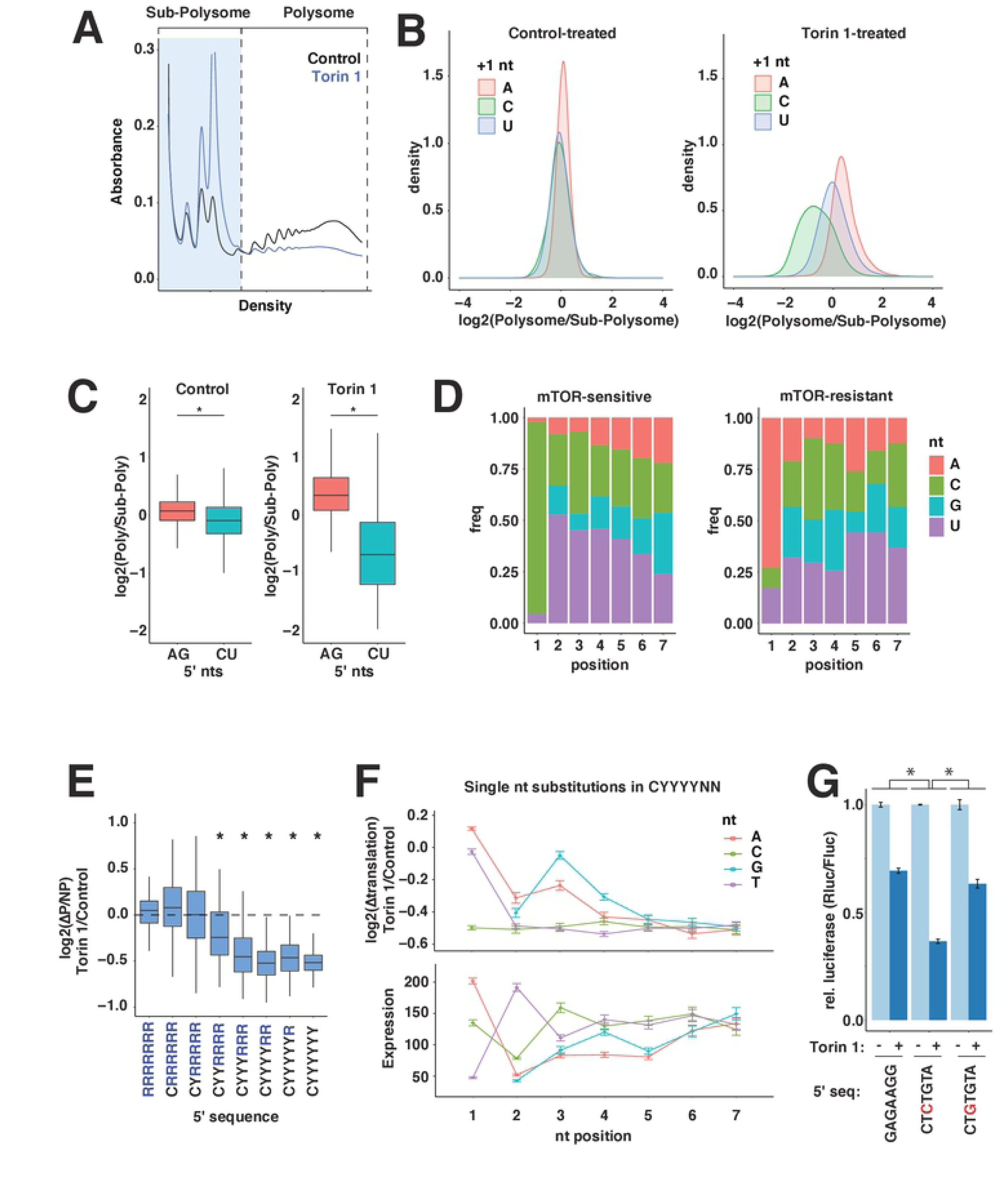
Initial nucleotides are determinants of translation rates. **(A)** Overview of 5pseq translation analysis. Extracts were prepared from HeLa cells stably expressing 5pseq library mRNAs and treated with vehicle (DMSO) or 250 nM Torin 1 for 2 h. Extracts were then centrifuged through 5-50% sucrose gradients and fractionated with constant monitoring of absorbance at 254 nm to obtain polysome profiles. RNA was extracted from the indicated polysome or sub-polysome fractions and used to prepare sequencing libraries. **(B)** Translation of 5pseq library mRNAs in control or Torin 1-treated conditions. Translation rates for library mRNAs isolated in (A) were estimated by Polysome/Sub-polysome ratios and depicted as density plots, separated by +1 nucleotide. **(C)** Box plots for translation rates of AG and CU mRNAs from (B). Significance by t-test. * p < 10^-10^. **(D)** 5′ nucleotide frequencies of mRNAs with mTOR-sensitive and mTOR-resistant translation. The frequencies of nucleotides at the first 7 positions of mRNAs with the 5% most and least mTOR-dependent change in translation from (B). **(E)** mTOR-sensitive translation increases with longer C/U 5′ sequences. Changes in translation following mTOR inhibition from (B) were determined for mRNAs with the indicated 5′ sequence motifs (R=A,G; Y=C,U). Significance by t-test comparison to mRNAs with R[7] 5′ sequences, * p < 0.01. **(F)** Sequence requirements of 5′ TOP motifs for translation and expression. The effect of individual nucleotide substitutions at the indicated 5′ positions in library mRNAs with 5′ CYYYYNN motifs on levels of mTOR-regulated translation from (B) and expression from Figure 2A. **(G)** The +3 C/U is key for TOP motif translation functions. Plasmids expressing mRNAs encoding Renilla luciferase and initiating with the indicated 5′ sequences were transfected into HEK-293T cells. Cells were incubated overnight, and then treated with vehicle (DMSO) or 250 nM Torin 1 for 2 h, and then analyzed for luciferase levels. Significance by two-way ANOVA, n=3. * p < 10^-5^.

This narrow range of translation rates may reflect that under basal conditions, the 5′ ends of most mRNAs are bound by the eIF4F initiation complex and effectively sequestered from other 5′ binding proteins [26]. In contrast, growth-repressive conditions destabilize eIF4F and expose mRNA 5′ ends to other translation regulators, such as 4EHP, Larp1 or decapping proteins. To test whether the translation of mRNAs becomes more sensitive to 5′ sequences under growth-repressing conditions, we again measured P/SP ratios in cells treated with the mTOR inhibitor Torin 1, which triggers growth arrest and inhibits eIF4F (Figure 4A) [27]. Under these conditions, the translation differences between mRNAs expanded to a significantly greater range, reaching 16-fold for transcripts differing by only a handful of nucleotides (Figure 4B). In particular, mRNAs with different +1 nucleotides experienced significantly different changes in translation. +1 A mRNAs maintained the highest levels of translation, while +1 C mRNAs were significantly repressed (Figure 4B). +1 U mRNAs were slightly repressed, but much less than +1 C mRNAs. Moreover, while the well-transcribed AG and CU mRNAs were similarly translated under growth-promoting conditions, the translation of CU mRNAs was selectively repressed by mTOR inhibition (Figure 4C).

### Functional requirements of TOP sequences

A +1 C is the defining feature of TOP mRNAs [28]. Indeed, the most mTOR-sensitive sequences within the 5pseq library closely resembled classical TOP motifs, with increasing enrichment of C/U nucleotides at positions close to the 5′ terminus (Figure 4D). mTOR-resistant sequences were primarily distinguished by a +1 A and no other obvious features (Figure 4D).

We showed previously that increasingly long series of C/U nucleotides within TOP motifs are correlated with greater repression [29]. Increasingly long series of C/U nucleotides also correlate with translation repression following mTOR inhibition in the 5pseq library (Figure 4E).

In this case, the maximum degree of suppression occurs at approximately 5 nt, which matches the number of nucleotides bound by the TOP suppressor Larp1 [8]. Our previous analysis of endogenous TSSs indicated a maximum sequence of 7 nt, but this likely reflects the fact that these TSSs are heterogenous, and longer C/U stretches ensure that a greater number of transcripts encode a maximal TOP motif. To systematically probe TOP motif requirements in the 5pseq library, we calculated the translation effect of varying each nucleotide in the canonical TOP sequence of CYYYYNN (Figure 4F). This confirmed the importance of the +1 C and the diminishing contribution of the next 4 nucleotides. Surprisingly, however, we found that the +3 position was particularly critical to TOP function. Replacement of a +3 C/U with a G was almost as disruptive as replacing the +1 C. In contrast, replacement of the +2 C/U with a purine was much less disruptive to translation regulation, but diminished expression level, as previously noted (Figure 2B). We confirmed these results with individual reporter constructs (Figure 4G).

### 5′ sequence link mRNA translation and stability

Because translation functions of 5′ sequences were most evident under growth-restricting conditions, we also wondered whether stability functions might be altered. To test this, we similarly measured library mRNA stabilities in cells with the mTOR inhibitor Torin 1 by blocking transcription with Actinomycin D (Figure 5A). Under these conditions, mRNA half-lives expanded to an approximately 4-fold range (compare Figure 5A to Figure 2D). +1 A mRNAs were the most unstable, while +1 C mRNAs were most stable, and +1 U mRNAs were in between (Figure 5A). We noticed that the stabilities appeared inversely related to their translation status, i.e. decreased translation correlated with increased stability. To determine the extent of this relationship, we compared the stability and translation of library mRNAs in both control and growth-inhibited conditions. Under control conditions, translation and stability were effectively uncorrelated (Figure 5B). Under growth-inhibited conditions, however, a strong correlation between these properties emerged, following a pattern that was again dominated by the identity of the +1 nucleotide (Figure 5B). +1 C mRNAs were simultaneously translationally repressed and stabilized, while +1 A mRNAs were subjected to the opposite regulation (Figure 5B). This argues that the translation and stability function of these mRNAs is linked by 5′ sequences.

**Figure 5.**
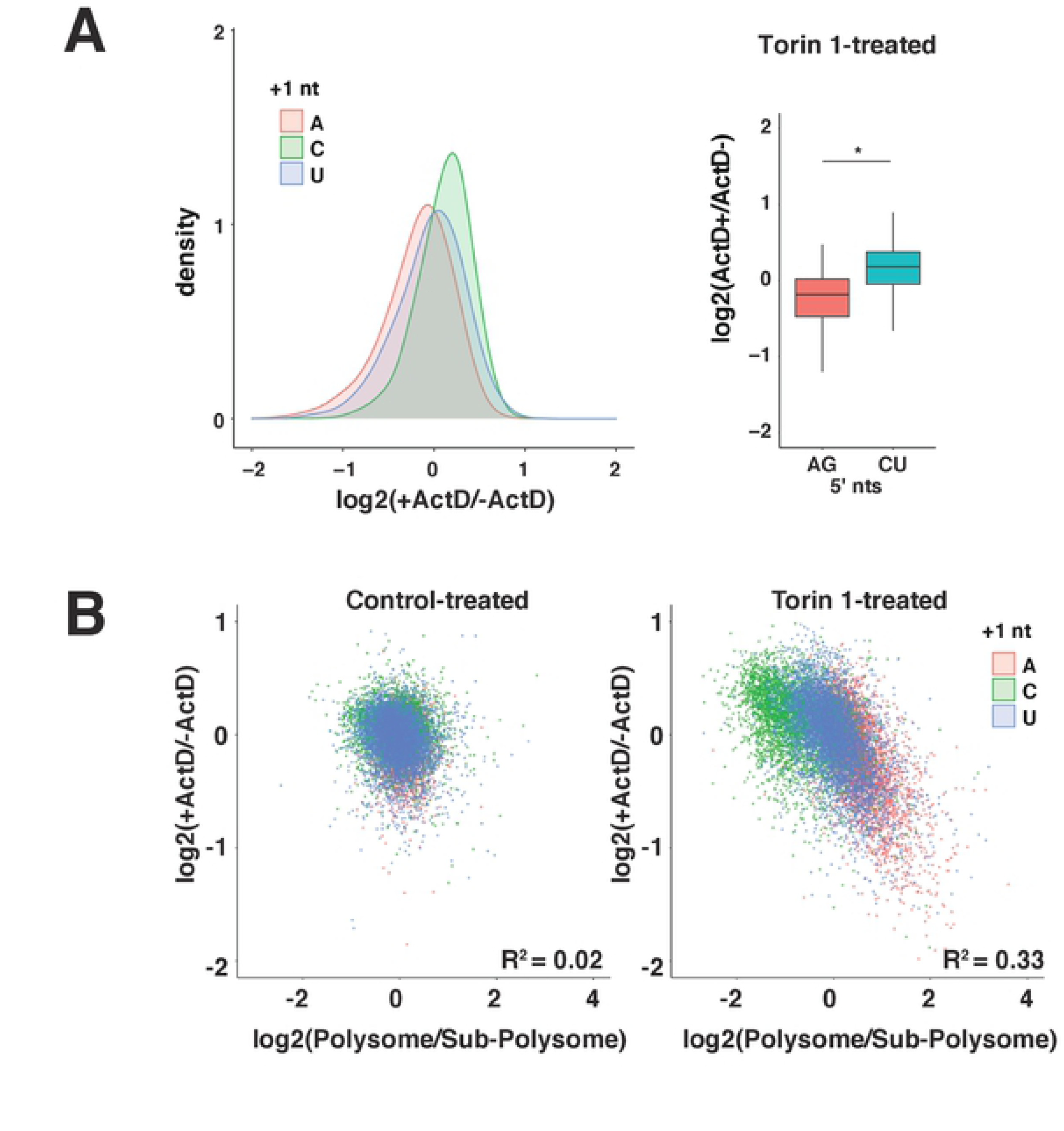
Translation and stability functions are correlated when cap-dependent translation is inhibited. **(A)** Large variation in library mRNA decay rates under mTOR-inhibited conditions. HeLa cells expressing library mRNAs were treated with 250 nM Torin 1 for 2 h, and then with 2 µg/mL Actinomycin D for 0 or 2 h in 2 biological replicates. Libraries prepared from extracted mRNA were analyzed to determine relative changes in levels between ActD-treated (ActD+) and untreated (ActD-) conditions. Left panel: Log2 changes in ActD+ vs ActD- levels are plotted separately for mRNAs initiating with the indicated nucleotide. Right panel: box plots of Log2(ActD+/ActD-) for mRNAs initiating with the indicated nucleotides. Significance by t-test for two biological replicates. * p < 10^-10^. **(B)** Comparison of decay and translation rates for library mRNAs under control and mTOR-inhibited conditions. Translation rates (Polysome/Sub-polysome) from Figure 4B and decay rates from (A) and Figure 2D for library mRNAs under mTOR-inhibited and control conditions. Datapoints are colored by +1 nucleotide.

### Endogenous mRNAs are optimized with TCT/TOP hybrids

The results described above show overlapping but distinct sequence requirements when comparing efficient transcription from TCT TSSs and growth-dependent translation/stability regulation of TOP mRNAs. Both systems require a +1 C for efficient transcription and translation regulation. However, the transcription system shows stronger preferences for a +2 T/U nucleotide while the translation system is particularly sensitive to the +3 nucleotide. A comparison of the expression level and translation function of 5′ 3 mers shows four specific 3- mers that optimize the function of both systems: CTC, CTT, CCT, and CCC (Figure 6A). To test whether these hybrid TCT/TOP sequences are enriched in endogenous mRNAs, we analyzed CAGE reads from several cell lines and tissues. Remarkably, these results showed that these 4 3mers were the most commonly used +1C sequences in nearly all of the datasets (Figure 6B). This argues that the TSSs of endogenous mRNAs reflect selection for both transcription efficiency and translation regulation. We noted one exception in liver 5′ sequences, where CTA replaced CCC for the fourth-most common 3 mer. CTA (as well as CTG) are both strong expression motifs, but only weakly sensitive to translation regulation. Interestingly, closer inspection of the liver TSS dataset revealed that the high frequency of CTA is primarily driven by the high expression of albumin, which primarily initiates with CTA. The selective pressures driving this sequence are not clear but may reflect other properties of TCT promoters that are optimized for consistent high expression.

**Figure 6.**
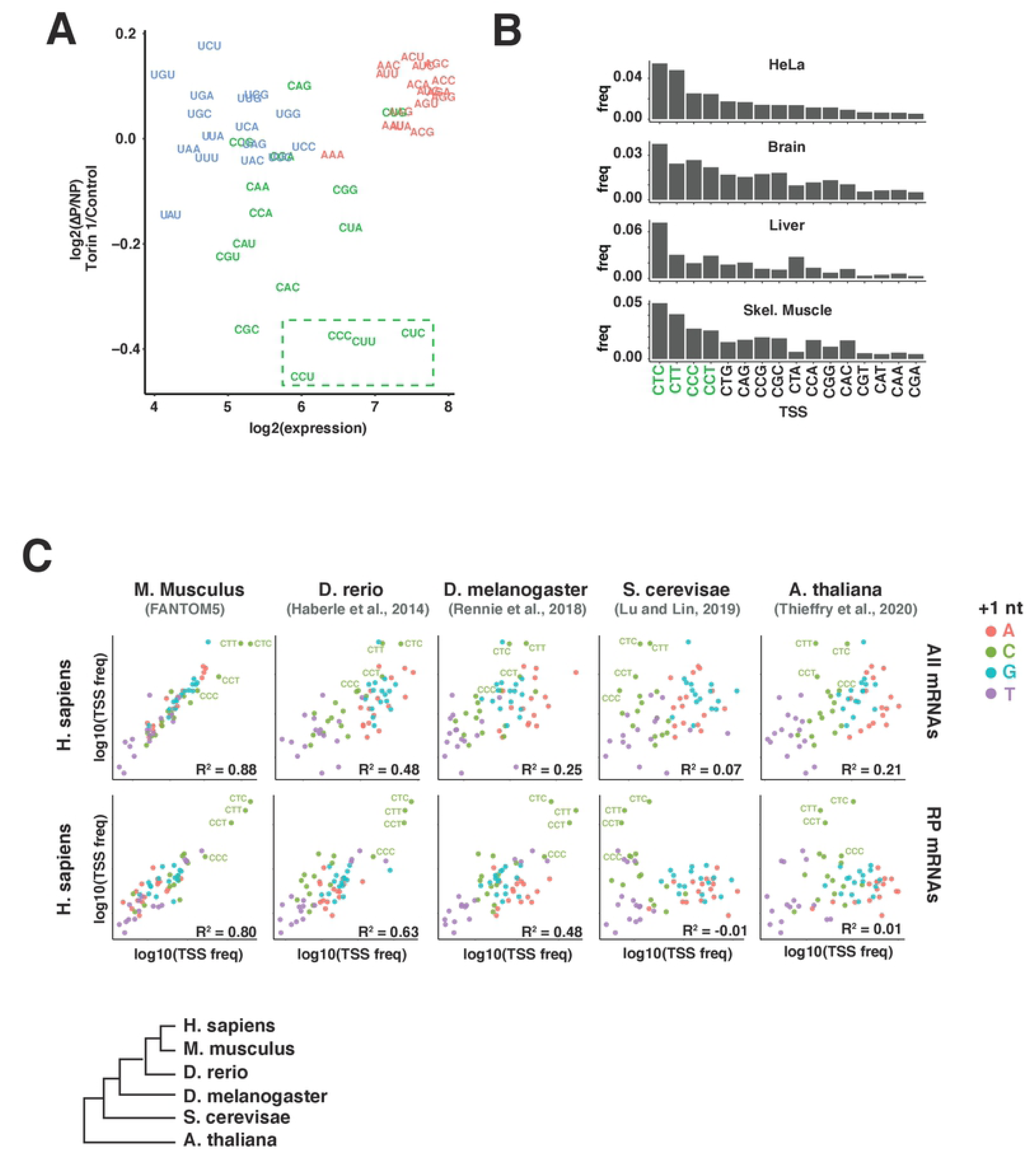
Endogenous TSSs are TCT/TOP hybrids optimized for both transcription and post-transcriptional regulation. **(A)** Comparison of translation and transcription functions of TSSs. The expression levels (Figure 2A) and mTOR-regulated translation (Figure 4B) of 5′ 3mer sequences are shown, colored by +1 nucleotide. The indicated TOP/TCT hybrids CTC, CCC, CTT, and CCT 5′ sequences maximize transcription and translation regulatory functions. **(B)** TCT/TOP 5′ sequences are enriched in endogenous mRNAs. Expression levels of the indicated +1 C 5′ sequences in hCAGE data from HeLa cells and the indicated human tissues. **(C)** Conservation of TSS usage across 6 species. Upper panel: Frequencies of 5′ 3mer sequences in CAGE reads mapping to 1 kb regions surrounding the 5′ ends of annotated transcripts in the indicated species were compared to 3mer frequencies in human (HeLa) hCAGE data. Lower panel: Frequencies of 5′ 3mer sequences in CAGE reads mapping to 1 kb regions surrounding the 5′ ends of annotated transcripts encoding ribosomal proteins from the indicated species. TCT/TOP sequences CTC, CCC, CTT and CCT are indicated.

### Evolutionary conservation of 5′ sequences

+1 C mRNAs are a unique class of transcripts that utilize specialized mechanisms for their transcription and post-transcriptional regulation. Aspects of this regulatory system have been reported in diverse species, including plants, flies, and throughout vertebrates. We therefore wondered how broadly patterns of 5′ sequences described here are conserved. To test this, we extracted reads aligned to promoter regions of annotated genes in CAGE datasets for 5 eukaryotic species (Figure 6C) [24, 30–33]. A limitation of this analysis is that different methods for TSS sequencing can affect the overall representation of specific sequences, even amongst datasets prepared with similar strategies. Nonetheless, we find that 5′ sequence usage is strikingly similar between humans, zebrafish, drosophila and even arabidoposis (Figure 6C). In contrast, there was much less similarity in TSS sequences between human and yeast datasets, which showed almost no expression of +1 C mRNAs (Figure 6C). Yeast TSSs, however, are similarly depleted of +1 T/U sequences.

In humans, the functional class most enriched for +1 C mRNAs are ribosomal protein (RP) genes, which almost all contain classical TOP motifs [28]. RP orthologues are highly conserved and so easily identified between species. To test whether this class of mRNAs utilizes the +1 C regulatory system across species, we compared RP gene TSSs between humans, zebrafish, flies, yeast and plants. 5′ sequence usage for RP genes in humans, zebrafish and drosophila were strikingly similar, preferring the same TOP/TCT hybrids that optimized expression and translation regulation in the 5pseq library (Figure 6C). These three species all express homologues of both the TOP binding translation regulator Larp1 and the TCT transcription factor TRF2, which is strong evidence that these regulatory systems function similarly across these species. Arabidopsis showed slight enrichment of +1C mRNAs in RP mRNAs, although the overall pattern of TSS usage is starkly different than observed in flies and vertebrates. This is consistent with a recent observation [34] that Larp1 targets a distinct subset of mRNAs in plants. Yeast TSSs showed no enrichment for any +1 C 5′ sequences and yeast does not express homologs of Larp1 or TRF2. Taken together, these results suggest +1 C regulatory systems emerged soon after the transition to multicellularity, but the specific evolutionary history remains unclear.

## Discussion

In this study, we systematically examined the functions of mRNA 5′ terminal sequences, a region uniquely positioned to influence all fundamental phases of the mRNA life cycle. Our results show that these sequences – and +1 nucleotides, in particular – define basic mRNA classes with functionally distinct patterns of transcription, translation and stability. We find that mRNAs initiating with AG or CU nucleotides are most efficiently transcribed. A +1 U is universally disfavored, consistent with its infrequency in endogenous mRNAs [35, 36]. Post-transcriptionally, +1 A and +1 C mRNAs are similarly well-translated and stable under growth-promoting conditions but differ significantly when growth signals are interrupted. +1 A mRNAs remain well-translated but unstable while +1 C mRNAs are translationally-repressed but stabilized. 5′ sequences beyond the +1 position, including variations in classical TOP motifs, can modulate these post-transcriptional functions. Importantly, the 5′ sequence patterns that are optimized for transcription and post-transcriptional control are present in endogenous mRNAs and broadly conserved across cell types and species.

At first glance, the different TSS patterns for +1 A and +1 C mRNAs – especially the strong preference of a +2 U for the latter – would seem to reflect the action of unique transcription mechanisms. Previous studies of endogenous +1 A and +1 C mRNAs have supported this hypothesis [13]. +1 A mRNAs are generally produced from Inr TSSs, while +1 C mRNAs are only known to be produced from TCT motifs [13]. Moreover, TCT and Inr TSSs were shown to recruit different configurations of the transcription machinery that are specialized for specific +1 nucleotides [17, 37]. In contrast, our system uses a single prosthetic CMV promoter that nonetheless efficiently yields both +1 A and +1 C mRNAs and recapitulates the TSS patterns that occur in endogenous mRNAs. How does a single CMV sequence recapitulate the specialized features of Inr and TCT motifs? One explanation is that the CMV promoter recruits multiple configurations of the transcription machinery for +1 C and +1 A mRNAs and positions them precisely at the same TSS location. A simpler explanation may be that 5′ AG and CU sequences, both in 5pseq library and endogenous mRNAs, reflect intrinsic preferences of Pol II for initiating transcription. Other defining characteristics of Inr and TCT-containing promoters are perhaps more important for determining the timing and levels of transcription.

Our results also reveal a striking global relationship between the stability and translation functions of 5′ sequences. Under growth-repressive conditions, +1 A mRNAs are globally well-translated but less stable, while +1 C mRNAs are translationally repressed but stabilized. For classical 5′ TOP sequences, this mechanism likely involves Larp1 [3, 38]. Our finding that that the same 5′ sequences necessary for translation regulation also impact stability implies that these mechanisms are tightly linked, at least within the context of the library mRNA used here. Unexpectedly, this inverse relationship between translation and stability under growth-repressive conditions extends across all 5pseq library sequences, not just TOP sequences (Figure 5B). Larp1 may generally recognize +1 C mRNAs, but it is unclear why +1 A mRNAs should behave similarly. One possibility is that these mRNAs are bound and stabilized by a Larp1-like protein that preferentially binds +1 A mRNAs, although we are currently unaware of any such protein.

A final question is to understand how the link between RNA translation and stability contributes to cellular function. Under growth-restrictive conditions, the link between translation and stability may offer two advantages. Many +1 C mRNAs, which include 5’TOP mRNAs, encode stable ‘housekeeping’ proteins, that are most needed during phases of cell growth. The simultaneous translation repression and stabilization of these mRNAs allows cells to temporarily (and rapidly) reduce protein production without forfeiting the investment in mRNA synthesis.

When permissive conditions return, cells are primed to resume production. Gentilella and colleagues recently proposed a similar “protective” model for Larp1 function that also involve the direct protection of small ribosomal subunits from degradation [39]. Additionally, the translation- stability link may be a mechanism for buffering the quantity of protein that is produced from each mRNA synthesized. In other words, a system that degrades mRNAs only when translated would define mRNA half-lives in terms of protein production rather than time, maintaining total protein production within a narrow range even amidst changing environmental conditions.

In summary, we have shown how different classes of 5′ sequences are linked to global patterns of transcription, translation and decay. This system provides a means for coordinating the expression of large classes of genes. Moreover, genes rarely initiate at a single TSS, yielding mRNAs with a spectrum of 5′ sequences. This allows cells to fine tune expression dynamics by producing mixtures of mRNAs with varying stabilities and translation in growth-promoting and inhibitory conditions. Importantly, these properties can be quickly changed by small shifts in TSS locations. Overall, the results described here are unlikely to have captured the full regulatory potential of 5′ sequences. In particular, we excluded +1 G 5′ sequences.

Although +1 Gs are common in endogenous mRNAs and follow expression patterns that are similar to +1 A mRNAs, specific +1 G 5′ sequences may possess unique functions. Second, our study only examined the first 7 nt. This allowed for a deeper sampling of all possible sequences, but excluded longer motifs that might encode functional (e.g. cap-proximal uORFs) or structural features that are significant for endogenous mRNA regulation. Further investigation will be necessary to answer these questions.

## Methods

### Materials

DMEM and TRIzol Reagent from Life Technologies; heat-inactivated FBS from Sigma Aldrich; T4 DNA Ligase I, polynucleotide kinase, proteinase K, Protoscript II reverse transcriptase, Phusion DNA polymerase, Vaccinia Capping System, T7 RNA polymerase from New England Biolabs; iTaq Universal SYBR Green Supermix and Bradford Protein Assay from Bio-rad; RNeasy Plus Mini Kit from Qiagen; DNA and RNA Clean and Concentrator 5 kits from Zymo Research; Endura Electrocompetent cells from Lucigen; Mighty Mix T4 DNA ligase from Takara; Dual luciferase reporter assay from Promega; and XtremeGENE 9 transfection reagent from Roche.

### Synthesis of 5pseq library

The 5pseq library plasmid was generated by introducing a Sal1 restriction site 25 nt downstream of the CMV promoter in the pCT3 plasmid (pCT3-TE2), a lentiviral plasmid based on pLJC1 (Addgene #87972) that encodes the mouse Eef2 5′ UTR and coding sequence for Renilla luciferase [9]. A DNA insert encoding a randomized 7 nt sequence adjacent to the CMV transcription start site was prepared using a two-step PCR amplification. First, primers TE117 and TE118 were used to amplify the promoter region of pCT3-TE2, while PCR of primers TE119 and TE120 was used to generate a dsDNA fragment containing the random 7 nt sequences flanked by part of the CMV promoter and part of the 5′ UTR. (Table 1). These were then combined in a second PCR reaction to generate a 420 nt fragment containing the entire CMV promoter, TSS, and partially overlapping the 5′ UTR. After clean-up, Gibson Assembly (NEB) was used to insert the dsDNA fragment into pCT3-TE2 that had been digested with NdeI and SalI. The ligated product was then electroporated into Endura^tm^ electro-competent cells (Lucigen #60242-1) in 2 separate reactions, grown in recovery media for 1 h, and then plated on agar plates with ampicillin. Dilutions were plated to estimate colony number. Plasmid was isolated from 17x 10^6^ colonies on 16 plates by scraping colonies into 50 mL tubes, and then isolating DNA by maxi-prep (Qiagen). To assess library complexity, the TSS/Promoter region was amplified by PCR using Illumina compatible primers (TE127 and TE111) from 5 ng plasmid using Phusion HF polymerase (NEB). Sequencing results were analyzed using custom Python scripts to quantify the frequency of each 7 nt TSS sequence.

**Table 1.**
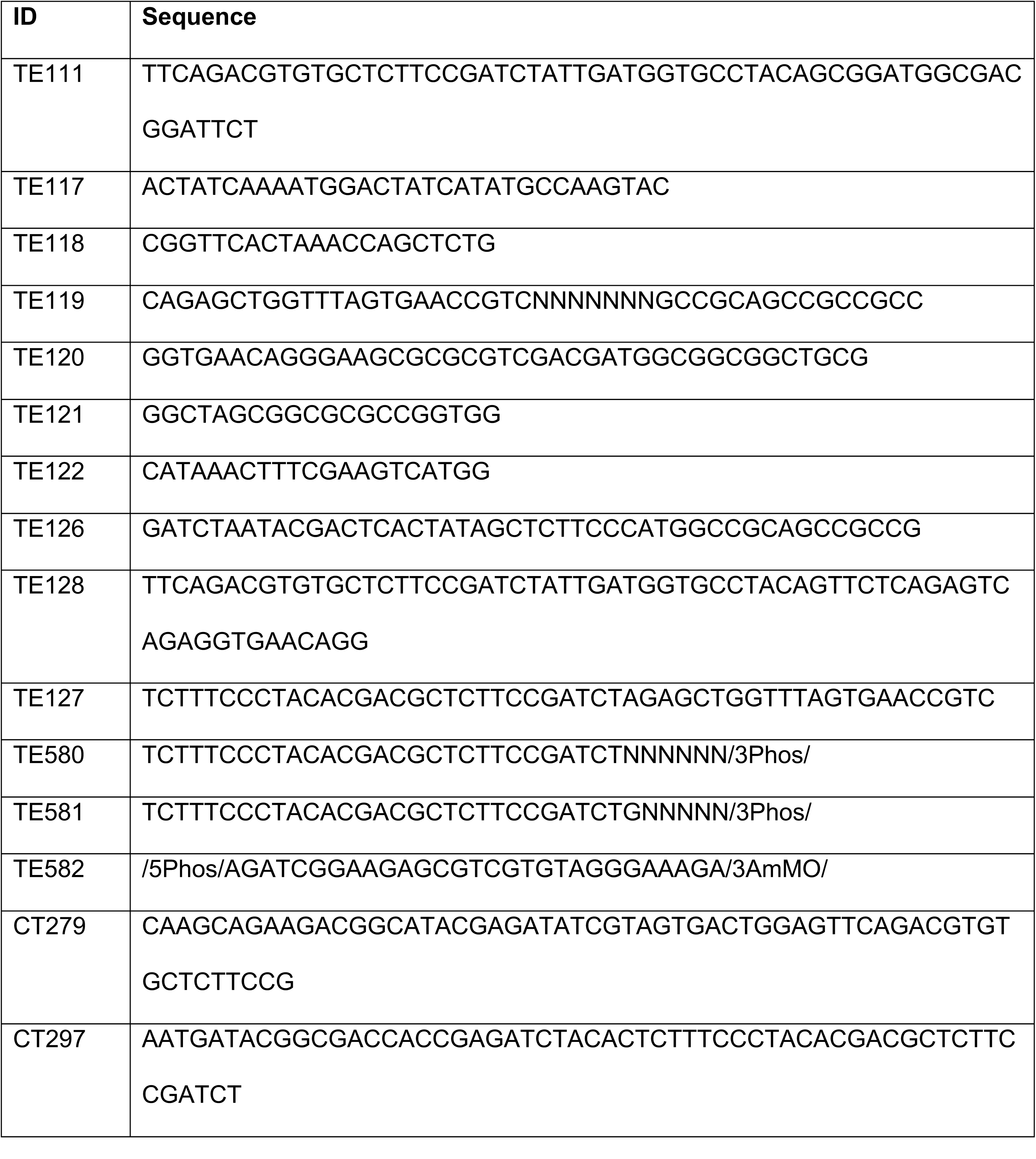
Primers used in library preparation and analysis.

### Viral infection of HeLa cells with 5pseq library

To prepare lentivirus for transducing the 5pseq library, HEK-293T cells were seeded on 3 15 cm plates at 13.5 million cells per plate and incubated overnight. The following day, cells on each plate were transfected with a mixture of 10 µg library plasmid, 9 µg psPax2 packaging plasmid (Addgene #12260), and 1 µg VSV-G envelope plasmid (Addgene #8454) using 100 µL PEI in 1 mL serum-free DMEM and incubated at room temperature for 10 min. Transfection mix was added drop-wise to cells. After 24 h media was replaced with 20 mL fresh DMEM + 10% FBS per plate, and cells were grown for an additional 24 h. To isolate virus, supernatant from cells was collected and centrifuged at 300 g for 5 min, and then filtered through a 0.45 µm filter to remove cellular debris. For infection, 14.5 mL virus was combined with 4.5 million HeLa cells in 15.5 mL fresh DMEM and 120 uL polybrene (2 mg/mL), and then seeded on a 15 cm plate.

After 24 h, media was replaced with 30 mL DMEM + 10% FBS supplemented with 0.4 µg/mL puromycin. After 48 h, media was replaced with fresh DMEM + 10% FBS. In the following days cells were trypsinized and seeded on plates for subsequent experiments. To analyze library expression, total RNA was extracted using TRIzol and used to prepare Illumina sequencing libraries as described below.

### Synthesis of capped spike-in mRNA

A DNA template for in vitro transcription was generated by PCR using oligos TE122 and TE126 on plasmid pCT-TE2 and Phusion DNA polymerase (NEB) in HF buffer (Table 1). PCR products were column purified (Qiaquick PCR purification kit) and 167 ng PCR product was used for *in vitro* transcription in a 100 ul reaction (30 mM Tris–HCl pH 8.1, 10 mM MgCl2, 2 mM spermidine, 0.01% Triton-X100, 10 mM DTT, 0.5 μl SuperaseIn (Ambion), 2 mM adenosine triphosphate (ATP), 2 mM guanosine triphosphate (GTP), 2 mM uridine triphosphate (UTP), 2 mM cytidine triphosphate (CTP), 2.5 μl T7 RNA polymerase (50 U/μl; NEB) at 37 °C for 2 hours, followed by 15 min DNAse I (NEB) treatment at 37°C. After clean-up (Zymo RNA Clean and Concentrator 5), 500 ng RNA was capped in a 20 µL reaction using Vaccinia capping system (NEB), following manufacturer’s instructions, followed by another round of clean up (Zymo RNA Clean and Concentrator 5). The final sequence of the spike-in mRNA is: m^7^G_CTCTTCCCATGGCCGCAGCCGCCGCCATCGTCGACGCGCGCTTCCCTGTTCACCTCTGACTCTGAGAATCCGTCGCCATCCGCCACCGGCGCGCCGCTAGCCACCATGACTTCGAAAGTTTATG.

### Sequencing library construction

The preparation of sequencing libraries from cells for quantifying 5pseq library expression is similar to preparing CAGE libraries for endogenous mRNAs, whereby a double-stranded splint adapter is ligated to the 3′ end of library cDNA [40].

The first step is to prepare the 3′ adapter. Two different adapter sequences were used, one ending with NNNNNN (TE580) and another ending with GNNNNN (TE581) to accommodate the frequent addition of a non-templated 3′ C during reverse transcription. Solutions of oligos TE580, TE581 and TE582 were prepared at 200 µg/mL in 1 mM Tris-HCl pH 7.5. Oligos TE580 and TE581 were each mixed at 2 µg/mL, separately, with oligo TE582 at 400 ng/mL of each oligo in 100 mM NaCl, denatured at 95 °C for 5 min, and then slowly cooled at 0.1 °C/s to 11 °C to anneal oligos. Annealed TE580/TE582 and TE581/TE582 were then combined at a 1:4 ratio and diluted to a final concentration of 200 ng/mL. Annealed adapters were then aliquoted and stored at -20 °C.

To prepare sequencing libraries, total RNA was reverse transcribed using a primer specific for library mRNA (TE121) and the Protoscript II reverse transcriptase (NEB). Following reverse transcription, RNA was hydrolyzed by the addition of 100 mM NaOH, heating to 98 °C for 20 min, and then pH neutralization by the addition of 100 mM HCl. cDNA was then cleaned up on silica columns (Zymo DNA Clean and Concentrator 5) and eluted in 6.5 µL of water. cDNA was then denatured at 65 °C for 5 min, and then placed on ice for 2 min. 1.5 µL of the adapter mixture (200 ng/mL) was prewarmed at 37 °C for 5 min then cooled on ice for 2 min, then combined with 6 µL cDNA and 15 µL Mighty Mix T4 DNA ligase reaction mix (Takara) and incubated overnight at 16 °C. cDNA was column-purified (Zymo DNA Clean and Concentrator 5) and eluted in 25.5 µL water.

PCR amplification of library mRNAs was performed in two steps. For the first step (PCR1), primers CT297 and TE128 were used to amplify library sequences from cDNA in a 50 µL reaction with Phusion polymerase in HF buffer (NEB) for 10 cycles. Amplified DNA was then isolated by column purification (Zymo DNA Clean and Concentrator 5) and eluted in 20 µL water. For the second step (PCR2), to determine the appropriate number of cycles, 2 µL of PCR1 product was used in three 20 µL reactions and amplified for 9, 11, or 13 cycles using oligos CT279 and CT297 with Phusion polymerase and HF reaction buffer (NEB). Each reaction was then analyzed by PAGE on a 12% TBE gel. The expected product is 203 nt. The number of cycles that yielded a single sharp band were used for the final PCR. For the final PCR, 2-8 µL of the initial PCR1 product, 5 µM each of the desired i5 and i7 Illumina dual index amplification primers, dNTPs, 1X HF reaction buffer and Phusion polymerase (NEB) were combined in a 80 µL reaction mix, then divided into 4 separate 20 µL reactions and amplified for the number of cycles determined in the test PCR. PCR products were combined, column-purified (Zymo DNA Clean and Concentrator), eluted in 15 µL water, and quantified using an Agilent Bioanalyzer. Library was then analyzed by Illumina sequencing on an Illumina NovaSeq or Hiseq 2500 analyzer.

### Quantification of library expression

To quantify the expression level of each 7 nt 5′ sequence, Illumina sequencing results were processed in several steps. First, the total counts of the spike-in mRNA, if used, were quantified using a custom Python script and removed from the FASTQ file. Second, each read was searched for a seed sequence present in the constant region of the library mRNA 5′ UTR (AGCCGCCGCC). Reads containing the seed sequence were processed to extract the first 30 nt of the mRNA sequence, including any non-templated G nts, and alignment position within the library plasmid. The frequencies of each 5′ sequence were then reported. Third, 5′ reads sharing a common “base” 5′ sequence but with varying numbers of non-templated G nts appended to the 5′ end were identified and grouped together. Non-template Gs were identified according to mismatches with the promoter region of the plasmid sequence. The counts for 5′ sequences within each group were summed to determine a final count for each common base sequence.

We note, this can only be determined for reads where the TSS begins with a non-G nt *<u>or</u>* aligns upstream of the +1 position of the 7 random nt sequence. The reason is that the TSS of any read initiating with a G that aligns within the random 7 nt sequence cannot be definitively distinguished from a read initiating downstream that has been extended by non-templated Gs. As a consequence, we considered only reads that definitively initiated at the +1 position of the random 7 nt sequence with a non-G nt.

### Analysis of 5pseq library translation

To measure translation rates of 5pseq library mRNAs, 13 million HeLa cells expressing the library were seeded on each of 4 15 cm plates in DMEM supplemented with 10% IFS and antibiotics and incubated overnight. Cells were then treated with vehicle (DMSO) or 250 nM Torin 1 for 2 h, and then washed 3 times in cold PBS- supplemented with 100 µg/mL cycloheximide, and then lysed in 1 mL polysome lysis buffer (20 mM Tris-HCl pH 7.4, 150 mM NaCl, 5 mM MgCl_2_, 1 mM DTT, 100 µg/mL cycloheximide, 1% Triton-X100). Cells were incubated for 5 min on ice, and then centrifuged 5 min at 14,000 rpm in a benchtop centrifuge to remove insoluble material. At this point, as a control, extracts from cells expressing a single classical TOP and non-TOP reporter mRNA, were added to library extracts. 300 µL of extract was then layered on top of a 5-50% sucrose gradient (20 mM Tris-HCl pH 7.4, 150 mM NaCl, 5 mM MgCl_2_, 1 mM DTT, 100 µg/mL cycloheximide, 5 or 50% sucrose) using a Biocomp GradientStation, and centrifuged at 36,000 rpm for 1.5 h in a SW41-TI rotor. Each gradient was then fractionated using a Biocomp GradientStation with constant monitoring at 254 nm separated into sub-polysome and polysome fractions. Fractions were supplemented with 0.5% SDS. Volumes were adjusted to 5.5 mL with water, then 10 ul capped spike-in (50 fg/ul) was added to each fraction, followed by digestion with 55 µL of proteinase K (NEB, 20 mg/mL) for 30 min at 50 °C. RNA was extracted with acid phenol, cleaned up with chloroform, and then precipitated with NaOAc and isopropanol. RNA resuspended in 11 µl water, of which 10 µl was used for library construction.

### Analysis of 5pseq library stability

To measure 5pseq library mRNA stability, 10 million library-expressing HeLa cells were seeded in each of 8 15 cm plates and incubated overnight. Cells were treated with vehicle (DMSO) or 250 nM Torin 1 for 2 h, and then treated with 2 µg/mL Actinomycin D for an additional 2 h or processed immediately. To extract RNA, cells were washed once in cold PBS, and then lysed in 2 mL Trizol containing 0.75 fg/µl capped spike-in. RNA was isolated according to the manufacturer’s instructions and resuspended in 21 µl water and quantified by UV absorbance, 10 µl of each sample was used to prepare Illumina-compatible libraries as described in the *Analysis of 5pseq Library Expression* section.

### Analysis of CAGE data

CAGE data was obtained from the sources listed in Table 1. To determine TSS frequencies, each dataset (Table 2) was aligned to the appropriate genome assembly (human: hg38, Mus musculus: mm10, Drosophila melanogaster: dm6, Danio rerio: Zv9, Saccharomyces cerevisiae: sacCer3/R64.2.1, Arabidopsis thaliana: tair10) using the STAR aligner [41]. Alignments were then retrieved for reads within a 1000 nt window centered on the TSSs of all annotated transcripts (human: GENCODE V38, Mus musculus: GENCODE VM23, Drosophila melanogaster: NCBI Refseq for dm6, Danio rerio: NCBI Refseq for Zv9, Saccharomyces cerevisiae: NCBI Refseq for sacCer3, Arabidopsis thaliana: TAIR10 genes) using SAMtools. The frequencies of all 7 nt TSS sequences in filtered reads were then determined using custom Python scripts. Ribosomal protein (RP) gene promoters for each species were identified based on gene name and manual curation. As with the transcriptome-wide TSS analysis, CAGE reads aligning to 1000 nt windows centered on the annotated TSSs for these transcripts were extracted using SAMtools and analyzed using custom Python scripts to quantify frequencies of 5′ 3-mers.

**Table 2.**
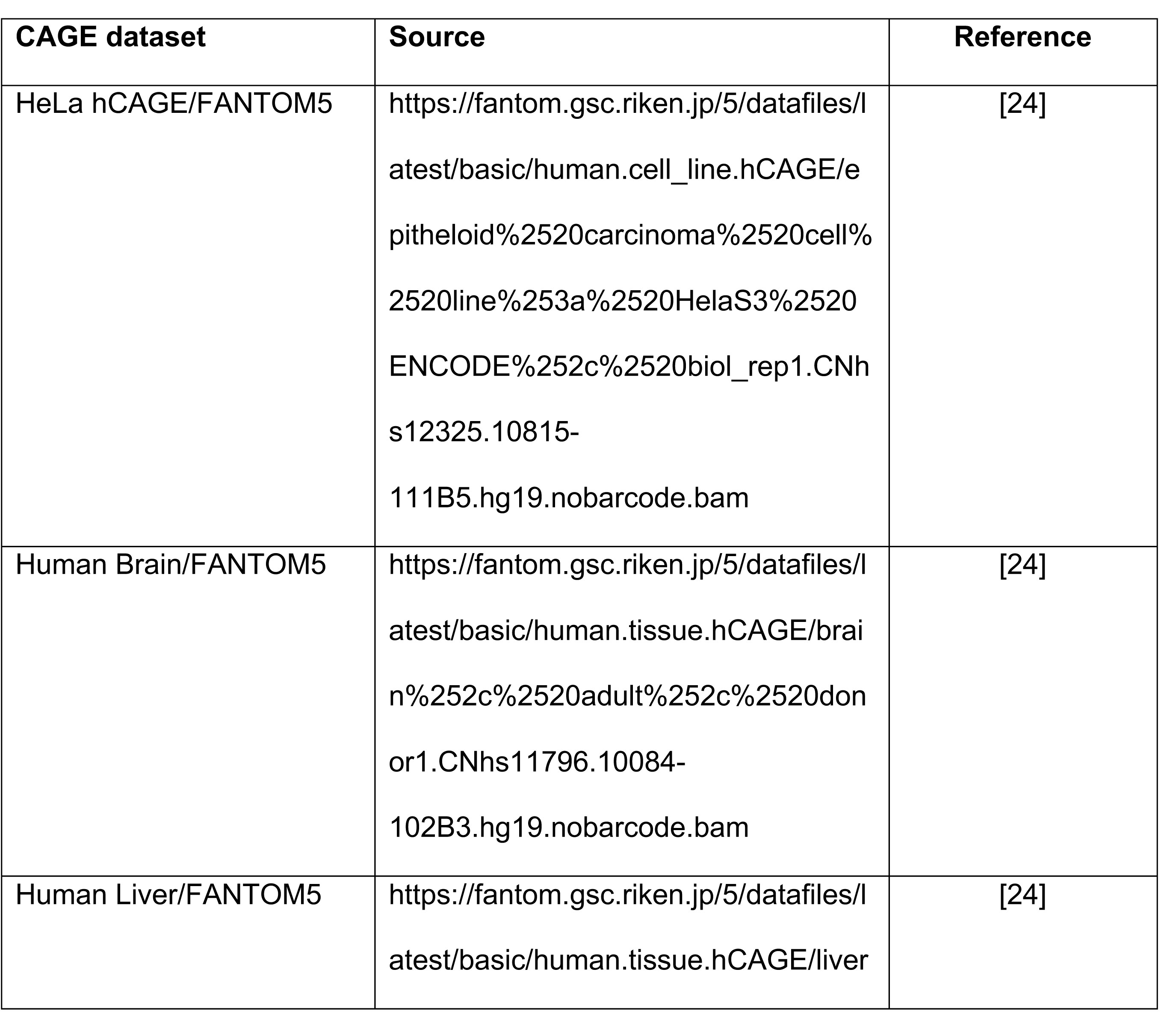
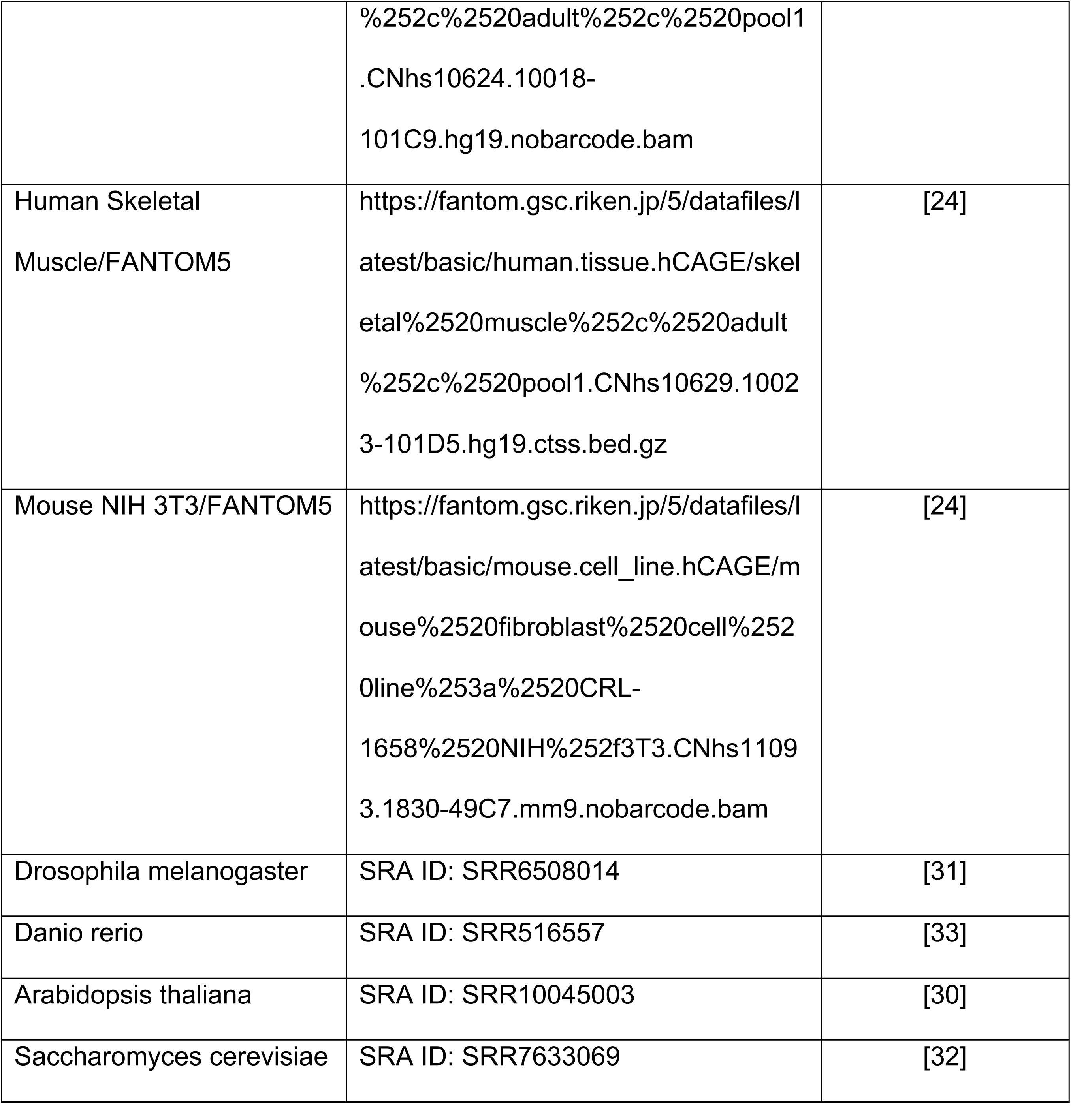
Sources for CAGE datasets.

### Translation and expression reporter assay

The indicated 5′ sequences were inserted into the library plasmid using Gibson assembly. HEK- 293T cells were transfected with 100 ng pIS0 (Addgene #12178, encoding firefly luciferase), 100 ng of the Renilla reporter and 800 ng of empty vector (1 µg total plasmid DNA) using XtremeGENE 9. After 24 h, cells were divided in 12-well plates at 0.3 million cells/well and incubated for an additional 24 h. For translation assays, cells were treated as indicated, and analyzed using the Promega Dual-Luciferase Reporter Assay System according to the manufacturer’s instructions. To measure expression, RNA was extracted using Trizol, reverse transcribed using Protoscript II (NEB), and quantified using qPCR with primers for renilla luciferase (forward: TCATGGCCTCGTGAAATCCCGT, reverse: GCATTGGAAAAGAATCCTGGGTCCG) and firefly luciferase (forward: GAGGCGAACTGTGTGTGAGA, reverse: GAGCCACCTGATAGCCTTTG). Levels of renilla luciferase were normalized to levels of firefly luciferase using the 1′1′Ct method [42].

## Data Availability

RNA-seq datasets have been deposited in the GEO Datasets database (GSE194192). Custom python scripts used in this manuscript are available upon request.

## Acknowledgements

We thank Mark Williams, Wendy Gilbert and Lucas Philippe for helpful discussions, and Michael Caplan for use of equipment and technical advice.

## Author Contributions

A.V. designed, synthesized, and analyzed 5pseq libraries. M.W. performed luciferase reporter experiments. A.V. and C.C.T. conducted bioinformatic analyses. C.C.T and A.V. designed all experiments and wrote the manuscript.

## Supplemental Data

Supplemental Figure 1. Nucleotide frequencies for the first 7 nt of plasmid and reporter 5′ terminal sequences. Sequencing libraries prepared from 5pseq plasmid or HeLa cells stably expressing the 5pseq library were analyzed to determine nucleotide frequencies in the first 7 nt of the expected (plasmid) or expressed (from cells) mRNA, respectively.

Supplemental Table 1. Expression levels of 5pseq mRNAs.

Read counts from the plasmid and reporter mRNA libraries under control conditions. Normalized expression levels are the mean reads per million (RPM) for reporter libraries divided by RPM for the plasmid library.

Supplemental Table 2. Stabilities of 5pseq mRNAs.

Columns are log2(fold change) (l2fc) and adjusted p-value (padj) calculated using DESeq2 for ActD-treated/untreated in control conditions in HeLa cells, ActD-treated/untreated in Torin 1- treated HeLa cells, and ActD-treated/untreated in control verus Torin 1-treated HeLa cells.

Supplemental Table 3. Translation rates of 5pseq mRNAs.

Columns are log2(fold change) (l2fc) and adjusted p-value (padj) calculated using DESeq2 for Polysome/Sub-polysome in control conditions in HeLa cells, Polysome/Sub-polysome in Torin 1-treated HeLa cells, and Polysome/Sub-polysome in control verus Torin 1-treated HeLa cells.

